# Distribution and abundance of bottlenose dolphin (*Tursiops truncatus*) over the French Mediterranean continental shelf

**DOI:** 10.1101/723569

**Authors:** Hélène Labach, Caroline Azzinari, Maxime Barbier, Cathy Cesarini, Boris Daniel, Léa David, Frank Dhermain, Nathalie Di-Méglio, Benjamin Guichard, Julie Jourdan, Nicolas Robert, Marine Roul, Nicolas Tomasi, Olivier Gimenez

## Abstract

The Mediterranean bottlenose dolphin (*Tursiops truncatus*) sub-population is listed as vulnerable by the International Union for Conservation of Nature. This species is strictly protected in France and the designation of Special Areas of Conservation (SAC) is required under the European Habitat Directive. However, little information is available about the structure, dynamic and distribution of the population in the French Mediterranean waters. We collected photo-identification data over the whole French Mediterranean continental shelf all year round between 2013 and 2015. We sighted 151 groups of bottlenose dolphins allowing the individually photo-identification of 1,060 animals. The encounter rate distribution showed the presence of bottlenose dolphins over the whole continental shelf all year round. Using capture-recapture methods, we estimated for the first time the size of the bottlenose dolphin resident population at 557 individuals (95% confidence interval: 216-872) along the French Mediterranean continental coast. Our results were used in support of the designation of a new dedicated SAC in the Gulf of Lion and provide a reference state for the bottlenose dolphin monitoring in the French Mediterranean waters in the context of the Marine Strategy Framework Directive.

## Introduction

The Common bottlenose dolphin (*Tursiops truncatus*, Montagu, 1821; hereafter bottlenose dolphin) is considered as a common species in the Mediterranean Sea. It has been observed along most of Mediterranean coast (Bearzi *et al.* 2009), preferentially over the continental shelf (Di Sciara *et al.*, 1993; Gannier, 2005; Gnone *et al.*, 2011), even though groups have also been observed offshore (Laran *et al.*, 2016). Both resident populations and transient individuals have been reported (Gnone *et al.*, 2011). Mediterranean bottlenose dolphins sup-population is genetically differentiated from populations inhabiting the contiguous eastern North Atlantic and the Black Sea and is structured into a Western and an Eastern population corresponding to habitat boundaries (Natoli *et al.* 2005).

The bottlenose dolphin Mediterranean sub-population is considered as “vulnerable” on the Red List of the International Union for Conservation of Nature (IUCN). It is listed in Annex II of the Washington Convention on International Trade in Endangered Species, in Appendix II of the Bern Convention for the Conservation of European Wildlife and Natural Habitats, in Appendix II of the Protocol to the Barcelona Convention on Specially Protected Areas of Mediterranean Importance (SPAMI) and is one of the only two species of cetaceans listed in Appendix II of the European Habitats Directive (92/43/CEE). It is also strictly protected in France by the decree of 1^rst^ July 2011 prohibiting, among other things, the destruction, capture and intentional disturbance of marine mammals. In addition, the bottlenose dolphin is the subject of a specific action plan under development by the Agreement on the Conservation of Cetaceans of the Black Sea, Mediterranean Sea and contiguous Atlantic Area (ACCOBAMS). In order to reach legal conservation objectives, the implementation of conservation strategies or action plans for a species requires the assessment of the population conservation status and the identification of trends in the population. Population indicators (e.g., distribution, abundance) should be regularly evaluated and compared with reference values through standardized long-term monitoring (Cairns *et al.* 1993; Dale & Beyeler, 2001).

In France, the monitoring program set up for the implementation of the European Marine Strategy Framework Directive (2008/56/EC; MSFD) recommends specific monitoring by photo-identification of resident coastal populations of marine mammal species, including bottlenose dolphins. Photo-identification is a methodology commonly used to monitor bottlenose dolphins (Shane *et al.*, 1986; Defran & Weller, 1999; Gnone *et al.*, 2011; Karczmarski & Cockcroft, 2014; Louis *et al.*, 2015)◻. Photo-identification allows individual monitoring for inferring population social structure, identifying movements and assessing population dynamics through the estimation of abundance and demographic parameters via capture-recapture (CR) methods (Hammond *et al.*, 2019; Hammond, 2009; Hammond *et al.*, 1990; Rosel *et al.*, 2011)◻.

In French Mediterranean waters, several studies on bottlenose dolphins have been conducted since the 1990s, mainly based on photo-identification (Bompar *et al.*, 1994; Dhermain *et al.*, 1999; Labach *et al.*, 2015; Labach *et al.*, 2011; Ripoll *et al.*, 2001), but remain local and punctual. The knowledge of the population’s structure, ecology and dynamic remain very poor and unequal.

In this study, we conducted the first large-scale survey of bottlenose dolphin based on photo-identification in the French Mediterranean waters. Standardized photo-identification data were collected all over the French Mediterranean continental shelf in each season over two years through a homogenized protocol by a network of organizations. The objectives of our study were to evaluate the distribution of bottlenose dolphin over the French continental shelf and to provide the first abundance estimate of the resident population.

## Methods

### Study area

The French Mediterranean waters present a great diversity and richness of habitats and seabed. The Gulf of Lion, from the Spanish border to Marseille, is a vast continental shelf limited to the north by a sandy and lagoon coastline and to the south by a broad slope cut by numerous canyons. The Corso-Liguro-Provençal basin (Riviera and west coast of Corsica) presents a rocky coastline prolonged by a very narrow continental shelf quickly giving way to an abrupt slope, cut by deep canyons, which debouches on the abyssal plain. To the east of Corsica, the reliefs are shallower with a larger continental shelf. The Corso-Liguro-Provençal basin and the Gulf of Lion are highly productive areas attracting a great diversity of species (D’ortenzio and Ribera Dalcaì, 2009)◻.

The study area covers the continental shelf of the French Mediterranean waters between the coast and the 500 m isobath, bounded by the Spanish border to the west, the Italian border to the east, and includes the whole Corsican coastline (Fig 1). The overall study area covers 24,481 km^2^ that was divided in three regions according to their geographic and topographic characteristics: Gulf of Lion (14,731 km^2^), Riviera (2,866 km^2^) and Corsica (6,884 km^2^).

**Figure 1:**
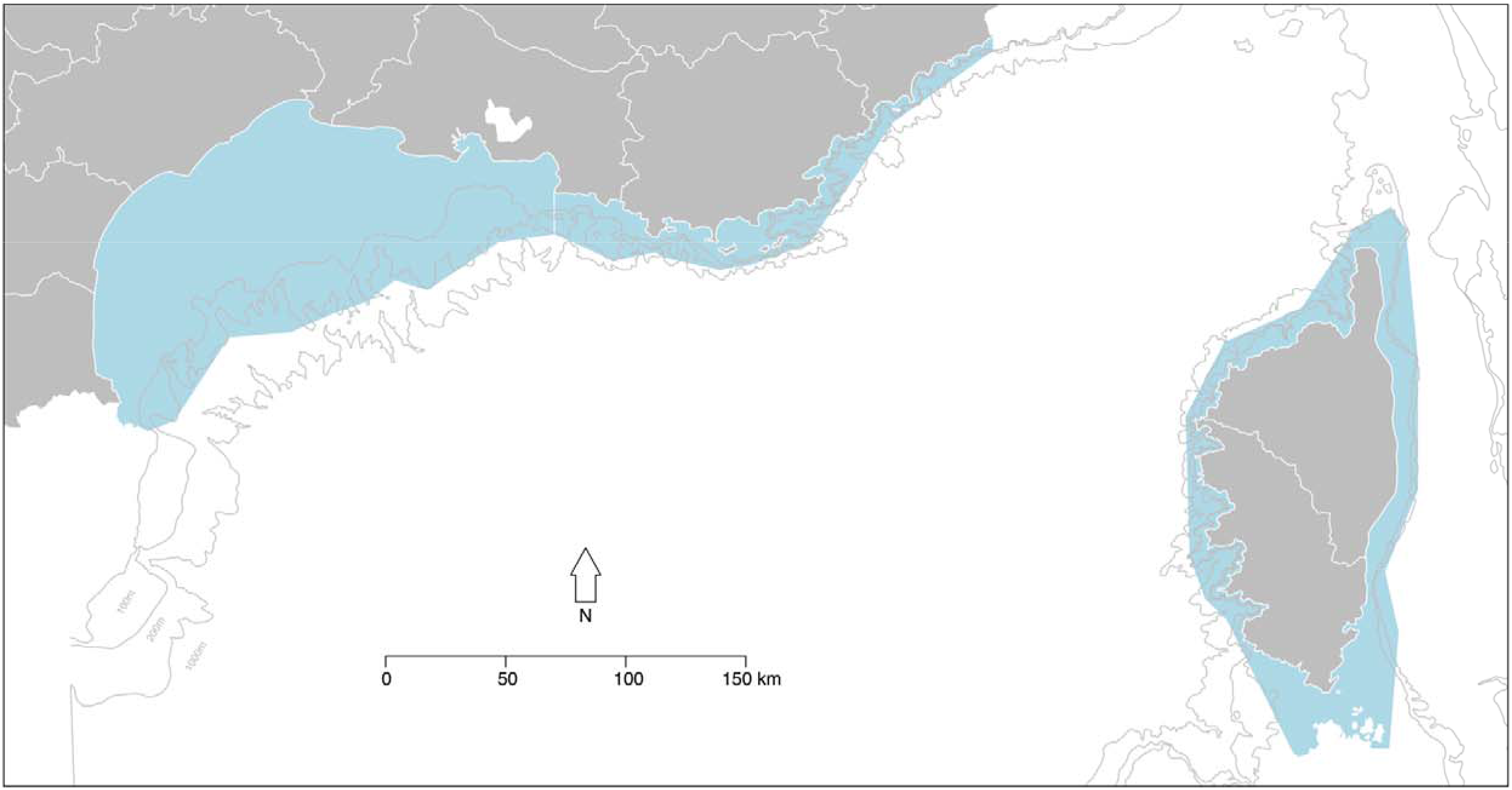
Study area (in blue) encompassing the French Mediterranean continental shelf in north-western Mediterranean Sea. The bathymetry is also displayed on the map.

### Data collection

To ensure a homogeneous sampling over the whole study area, each region was divided in sub-regions of similar area (4 in Gulf of Lion, 2 in Riviera and 3 in Corsica) and assigned to 5 local structures involved in marine mammals monitoring (BREACH, CARI, EcoOcean Institut, GECEM and Parc naturel regional de Corse). Each partner conducted 4 days of boat-based survey in good weather conditions in each season during 2 years in the sub-regions assigned to it. We carried out surveys between summer 2013 and summer 2015 using small sailing and motor boats. We designed these surveys to locate and photo-identify bottlenose dolphins and optimize the study area’s sampling coverage. All partners applied a standard common protocol using a digital application for the data collection specifically designed with Cybertracker (https://www.cybertracker.org/), systematically recording survey tracks with a GPS receiver. When we encountered a group of bottlenose dolphins, we recorded the position of first contact, group size and composition along with group main activity. Whenever possible, we took pictures of both sides of dorsal fins of all individuals of the group with digital reflex camera. We gathered all data and best pictures of each sighting in a common database and uploaded the data on the international web database INTERCET (http://www.intercet.it/).

### Photo-identification

We identified individuals using natural marks: scars, nicks, and scratches on their dorsal fins (Würsig and Jefferson, 1990; Würsig and Würsig, 1977)◻. We selected best quality pictures (methodology described below in *Abundance estimation* paragraph) of both profiles of each individual for each sighting and created catalogs of dolphins identified with the history of their sightings. Each partner compared its own catalog with all the others, hence leading to three regional catalogs and one global catalog containing the encounter history of each dolphin photo-identified during the study period.

### Survey effort

We defined the survey effort as the length (in km) of track actively traveled prospecting the area with naked eyes by three observers in favorable weather conditions (wind speed lower than Beaufort 3 and good visibility).

### Group size

We defined a group as all the dolphins seen with naked eyes during the sighting. The estimated group size is the estimated number of individuals observed or photo-identified whenever the latter figure is greater than the estimated one.

### Distribution

We calculated the encounter rate (ER) as the number of sightings per km of effort traveled in each region and within each 5’×5’ cell of the Marsden grid WGS 84. All maps and spatial analyses were done in R 3.5.0 (R Core Team, 2018)◻.

### Abundance estimation

To estimate the abundance of bottlenose dolphins occurring within the study area, we fitted CR models to the photo-identification data (Hammond *et al.*, 1990)◻. We defined a capture as the time an individual was identified with photo-identification, and a recapture as the resighting of an individual already seen during the project.

We scored best pictures of each dolphin sighting according to their quality and the distinctiveness of animals using 1 for good, 2 for medium and 3 for bad (Ingram, 2000)◻. We used only medium and good quality photos (quality scores = 1 or 2) of moderately and well-marked individuals (distinctiveness score = 1 or 2).

Because mortality most likely occurred during the study period, we used the Cormack-Jolly-Seber (CJS) (Cormack, 1964; Jolly, 1965; Seber, 1965)◻ model to estimate abundance while accounting for apparent survival (the product of true survival and fidelity) and a recapture probability less than one. We considered the eight seasons as our capture occasions. The main assumptions underlying the CJS model (Lebreton *et al.*, 1992) are 1) the population is demographically open (i.e. natality and mortality events occur) during the study period; 2) all individuals are correctly identified at each capture occasion and 3) the marks are considered permanent. Although these assumptions were unlikely to be violated in our study, we formally evaluated the quality of fit the CJS model to the data at hand (see next paragraph).

We performed three distinct analyses corresponding to the sightings made in the Gulf of Lion, the Riviera and along the continental coast (Gulf of Lion plus Riviera). We did not pursue CR analyses with the Corsican sightings because of the insufficient number of recaptures (Table 1). To fit CR models, we used the RMark package (Laake and Rexstad, 2008) which calls the MARK program (White and Burnham, 1999) in program R. We use the R package R2ucare (Gimenez *et al.*, 2018) to assess the quality of fit of the CJS model to data (Pradel *et al.*, 2005)◻. While trap-dependence was not detected, we detected a transient effect that we accounted for by using a two-age class for survival (Roger Pradel *et al.*, 1997). Individuals that were sighted only once were part of the first age-class (transients were included in this class) while all the others were part of the second. The age in CR analysis was considered as the time passed since the animal was first sighted (Madon *et al.*, 2012)◻. The proportion of transients was estimated and the abundance estimate amended accordingly (Madon *et al.*, 2012)◻. To test and account for the presence of heterogeneity in the detection probability, we used CR mixture models (Pledger *et al.*, 2010) in which animals belong to different classes of detection in proportions to be estimated (Gimenez *et al.*, 2017). For each analysis, we fitted four models incorporating a season and/or heterogeneity in the recapture probability while survival was considered constant over time. To determine the most parsimonious model, the model with the lowest AICc score (Akaike Information Criterion corrected for small sample sizes) (Burnham and Anderson, 2002) was selected (Appendix 1). The selected model was then used in a non-parametric bootstrap procedure (with 500 iterations) to calculate 95% confidence interval for population size (Cubaynes *et al.*, 2010)◻.

**Table 1:** Sightings and photo-identification of bottlenose dolphins

|  | Sightings | ER | Group size<br>(SD) | Right<br>profiles | Left<br>profiles | Captures | Recaptures | Identified<br>individuals | Recaptured<br>individuals |
| --- | --- | --- | --- | --- | --- | --- | --- | --- | --- |
| Corsica | 41 | 0.012 | 5.3 (4.5) | 140 | 130 | 167 | 35 (21%) | 132 | 26 (20%) |
| Riviera | 18 | 0.003 | 15.7 (10.3) | 227 | 207 | 260 | 113 (43%) | 147 | 45 (31%) |
| Gulf of |  |  |  |  |  |  |  | 834 | 248 (30%) |
| Lion | 92 | 0.007 | 16.6 (13.2) | 920 | 895 | 1278 | 446 (35%) |  |  |
| Global | 151 | 0.007 | 13.6 (12.5) | 1287 | 1,232 | 1705 | 648 (38%) | 1060 | 334 (32%) |

Number of sightings, encounter rates (ER), mean group size and standard deviation (SD), number of right and left profiles pictures, number of captures and number and percentage of recaptures, number of individuals identified, number and percentage of individuals captured more than once.

Because we used only well and moderately marked individuals (assumed to be adults) in the CR analyses, the total abundance including poorly marked individuals (juveniles and neonates) was obtained by correcting the CR-estimated abundance by the proportion of poorly marked individuals (Williams *et al.*, 1993)◻.

## Results

### Survey effort

We traveled a total of 21,464 km in survey effort. The distribution of the effort between the 3 regions was heterogeneous with a high coverage of Riviera but low coverage of Corsica and the offshore areas of Gulf of Lion. Summer was the best prospected season, autumn and winter being less prospected in the three regions (Fig 2).

**Figure 2:**
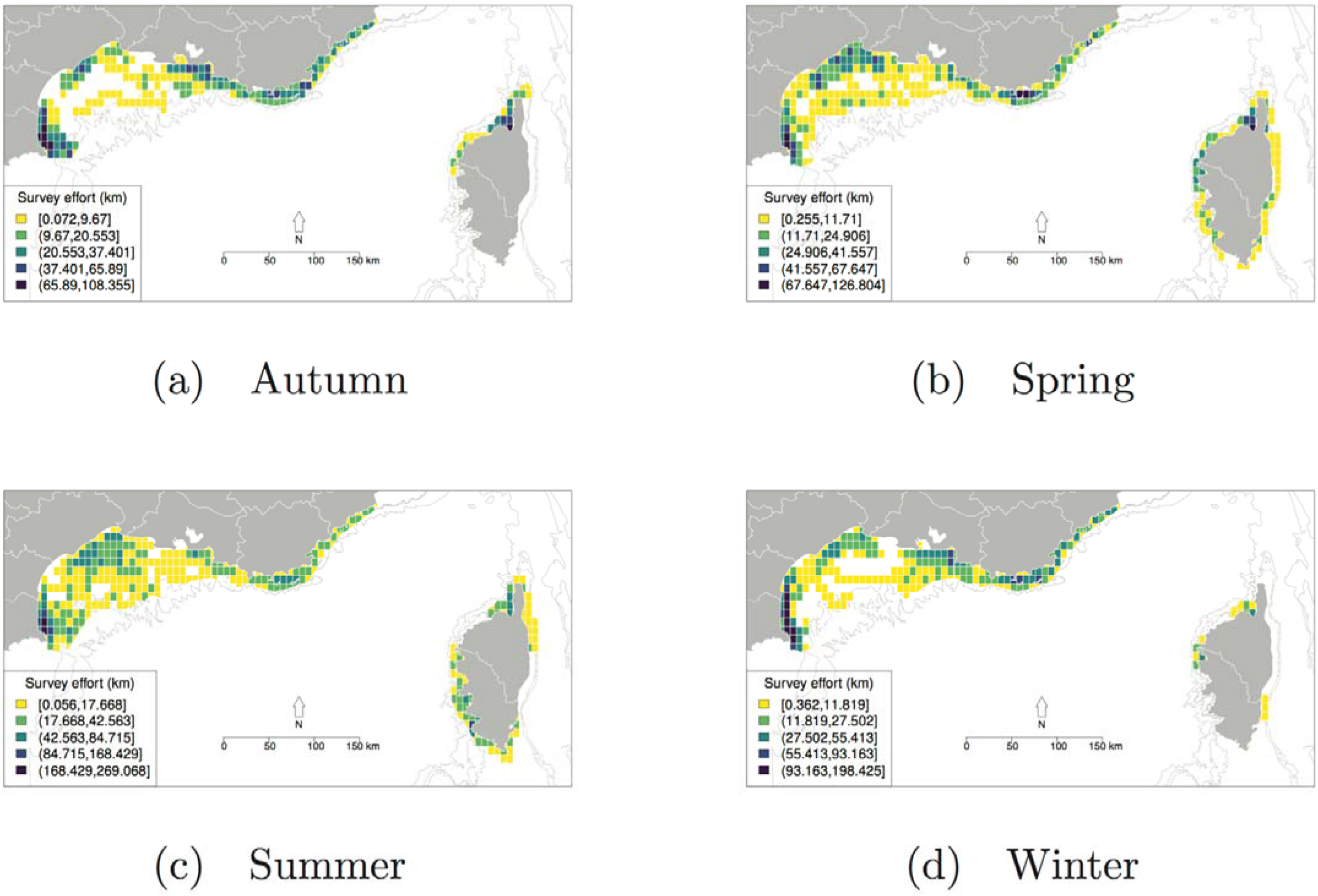
Seasonal distribution of survey effort (number of kilometers actively traveled per 5’x5’ cell) between 2013 and 2015 over the French Mediterranean continental shelf.

### Sightings and photo-identification

We sighted 151 groups of bottlenose dolphins during the project. Group size was highly variable in the three regions, mean group size was similar in Riviera and Gulf of Lion and lower in Corsica (Table 1).

We made a total of 1,705 photo-identifications of 1,060 dolphins (Table 1), of which 32% were observed more than once during the project. The percentage of individuals recaptured was higher in Riviera and lower in Corsica. We did not record any recapture between continental and Corsican coast during the project, while we observed 53 individuals in both Riviera and Gulf of Lion.

### Distribution

We sighted bottlenose dolphins in the whole study area all year round (Fig. 3). Global ER was higher in Corsica and lower in Riviera (Table 1). In Riviera, ER was higher in spring, while in Gulf of Lion and Corsica, ER was higher in summer.

**Figure 3:**
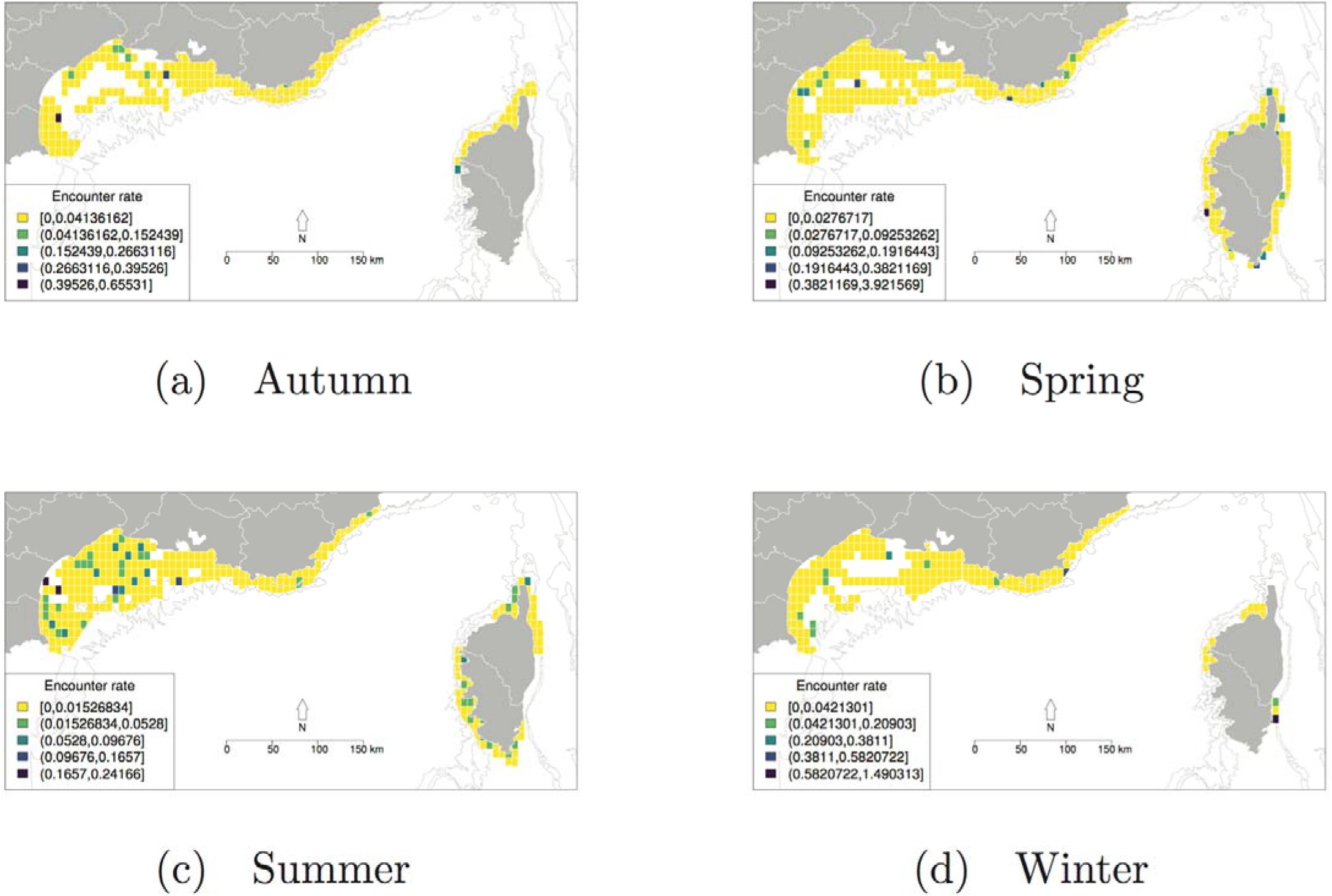
Seasonal distribution of bottlenose dolphins over French Mediterranean waters between 2013 and 2015. Encounter rates (number of sightings/km) per 5’x5’ cell.

### Abundance estimates

We excluded 15% of the pictures from the analyses because of their low quality (score 3). The percentage of moderate and well-marked individuals was 59% in Riviera, 77% in Gulf of Lion and 76% in the whole continental coast. Many dolphins (68% in continental coast) were seen only once. The maximum number of captures was 6 for two dolphins (Table 2).

**Table 2.** Distribution of individuals per number of captures

|  | 1 | 2 | 3 | 4 | 5 | 6 | Total |
| --- | --- | --- | --- | --- | --- | --- | --- |
| Riviera | 79 | 9 | 5 | 3 | 1 | 0 | 97 |
| Gulf of Lion | 411 | 100 | 51 | 15 | 1 | 2 | 580 |
| Continental coast | 458 | 123 | 61 | 21 | 6 | 2 | 671 |

Number of dolphins identified 1, 2, 3, etc. times in each dataset.

**Table 3:** Abundance estimates (N) and 95% confidence intervals in Riviera, Gulf of Lion and Continental coast in each season. For Winter and Summer 2014 in Riviera, the recapture probabilities were estimated very low, which impeded the estimation of abundance.

|  |  | Summer | Autumn | Winter | Spring | Summer | Autumn | Winter |
| --- | --- | --- | --- | --- | --- | --- | --- | --- |
|  |  | 2013 | 2013 | 2013-2014 | 2014 | 2014 | 2014 | 2014-2015 |
| Riviera | N | 57.74 | 33.86 | NA | 57.78 | NA | 52.63 | 58.39 |
|  | 2.5% | 25.52 | 18.85 | NA | 21.56 | NA | 22.65 | 23.17 |
|  | 97.5% | 98.45 | 54.12 | NA | 151.24 | NA | 128.96 | 113.43 |
| Gulf of<br>Lion | N | 377.36 | 297.87 | 539.56 | 338.44 | 558.28 | 499.50 | 494.55 |
|  | 2.5% | 124.21 | 147.82 | 266.82 | 236.38 | 453.18 | 399.50 | 412.48 |
|  | 97.50% | 875.67 | 631.09 | 1395.10 | 481.26 | 679.21 | 596.49 | 604.17 |
| Continental<br>coast | N | 199.47 | 307.38 | 888.72 | 446.28 | 775.44 | 646.10 | 635.20 |
|  | 2.5% | 134.17 | 221.38 | 491.81 | 349.91 | 613.78 | 520.81 | 516.36 |
|  | 97.5% | 297.51 | 460.92 | 1717.56 | 535.68 | 963.60 | 788.23 | 748.54 |

According to AICc values (Appendix 1), the model best supported by the three datasets was the model considering two age classes in survival and season-dependent recapture probabilities. The mean ratio of transient animals was estimated to 0.69 (95% CI 0.36-0.85) in Riviera, 0.45 (95% CI 0.37-0.53) in Gulf of Lion and 0.41 (95% CI 0.33-0.50) in continental coast.

Mean total abundance (corrected by the ratio of moderately and well-marked individuals) of resident population has been estimated at 43 (95% CI 4-58) individuals in Riviera, 444 (95% CI 304-555) in Gulf of Lion and 557 (95% CI 216-872) along the continental coast.

## Discussion

Our study provides the first large-scale dedicated photo-identification survey for the bottlenose dolphin in the French Mediterranean waters. We demonstrate the power of a collaborative and coordinated survey to study a mobile species at the scale of a population. Our results show that the whole continental shelf is frequented by bottlenose dolphins, including the entire Gulf of Lion, all year round. We also confirmed the presence of a resident population along the French Mediterranean coasts, for which we provided the first abundance estimate in Riviera and Gulf of Lion.

The prospecting effort of 21,464 km covered 87% of the study area. We found heterogeneity in this effort between the three regions which we explained by a later start of the survey in Corsica and more difficult survey conditions in the Gulf of Lion because of the important offshore area which demands long-distance offshore survey trips. The entire coastline of the French Mediterranean is often subject to difficult weather conditions limiting survey effort, especially in Winter.

The global encounter rate (0.007) was higher than the encounter rates (0.0041 with CV = 0.17 in winter and 0.0028 with CV = 0.2 in summer) obtained with the program “Surveillance Aérienne de la Mégafaune Marine” (SAMM), a comprehensive aerial survey of marine megafauna conducted by the French Biodiversity Agency in 2011 and 2012 over the whole French Exclusive Economic Zone (EEZ), encompassing continental shelf, slope and oceanic waters (Laran *et al.*, 2016)◻. The ER in Riviera (0.003) and in Corsica (0.012) was also higher than the maximum ER obtained by (Gnone *et al.*, 2011)◻ between 1994 and 2007 in Provence (ER = 0.0006) and in Corsica (ER = 0.0086), which might be due to an increase in dolphin abundance in these two regions, but the different time scale and different sampling methods make the comparison difficult.

The distribution of ER showed that bottlenose dolphins were present over the entire French Mediterranean continental shelf all year round. The higher ER in summer in Gulf of Lion and Corsica was consistent with the results of the SAMM survey, which showed higher ER in winter than in summer in the global EEZ, but a distribution more important in offshore waters in winter and in coastal waters of Gulf of Lion and Corsica in summer (Laran *et al.*, 2016)◻. These results, together with the detection of a strong transient effect in the CR analyses, suggest a seasonal migration of bottlenose dolphins between offshore waters in winter to coastal waters in summer, especially in Gulf of Lion and Corsica. The sighting of 53 dolphins both in Riviera and Gulf of Lion also points towards eastward and westward movements. No movement between the continental areas and Corsica was observed during the project, although 5 individuals were identified both in Corsica and along continental coast in previous studies (Gnone *et al.*, 2011). The identification of distinct units and the characterization of connections between them is the object of ongoing work using population genetic and social structure analyses based on photo-identification and biopsy data collected during the present study. The higher percentage of badly marked individuals (41%) suggests, in Riviera, a higher percentage of immature dolphins than in Gulf of Lion (23%).

The robust estimation of abundance relies on the validation of CR model assumptions. The two-year sampling period and the fact that newborns were observed in the study area suggest that assumption 1 of the CJS model is likely to have been respected. Assumptions 2 and 3 are ensured by the fact that only moderately and well-marked individuals with good-quality photographs were included in the analysis. Also, if the marks evolve, the short sampling period would allow to recognize the animals.

The average total population along the continental coast between 2013 and 2015 estimated at 557 (95% CI 216-872) individuals was higher than the estimates of the only previous census campaign dedicated to bottlenose dolphins in the same area, which estimated by observed count (ignoring imperfect detection), the number of bottlenose dolphins between 200 and 209 in the Gulf of Lion and 16 in Provence (Ripoll *et al.*, 2001)◻. These figures are not inconsistent with our abundance estimates which were corrected to account for imperfect detection. Our abundance estimates are coherent with the results obtained from the program SAMM with the distance sampling methodology, which estimated the absolute abundance of bottlenose dolphins in French territorial water (within 12 miles of the coast) at 350 (95% CI 150-900) dolphins inside the Pelagos Sanctuary and 500 (95% CI 115-2,500) outside in Winter and at 1,800 (95% CI 900-3,500) individuals inside the Pelagos Sanctuary and 450 (95% CI 120-1,700) outside in Summer (Laran *et al.*, 2016)◻.

### Implications for conservation

Our study provides an operational framework as well as a reference state for the implementation of a long-term large-scale monitoring of the resident bottlenose dolphin population in the French Mediterranean waters in the framework of the Marine Strategy Framework Directive. We shared the data on the international webGIS platform INTERCET (http://www.intercet.it/) which will allow to enlarge the study of this species beyond French boundaries to the basin and Mediterranean scale.

The results of our study together with those from the SAMM survey (Laran *et al.*, 2016)◻ led to an update of the Mediterranean bottlenose conservation status in the national IUCN Red List which was changed from “vulnerable” in 2009 to “nearly threatened” in 2017 because of knowledge improvement. Our demonstration of the presence of bottlenose dolphins in the entire Gulf of Lion led France to submit the designation of a dedicated offshore SAC encompassing the whole Gulf of Lion continental shelf beyond the territorial waters and to the recognition of this area as an important marine mammal area (IMMA) for bottlenose dolphins (https://www.marinemammalhabitat.org/imma-eatlas/). Our results will also contribute to update the ACCOBAMS bottlenose dolphin conservation plan.

We recommend that the photo-identification monitoring of bottlenose dolphins over the French Mediterranean continental shelf is continued in the long term to allow the identification of trends in the population and the implementation of adaptive management of the species at the sub-regional scale.

## Acknowledgements

We acknowledge MAVA Foundation, the French Biodiversity Agency and the Pelagos Sanctuary, who financially supported the GDEGeM project and also thank Fondation de France for funding through the INTERACT project. We thank all the partners for their involvement in the data collection: BREACH, CARI Corse, EcoOcéan Institut, GECEM and Parc naturel régional de Corse as well as all the people who participated to the survey. We also acknowledge Guido Gnone and Michela Bellingeri for their support in the use of the INTERCET platform.

## Appendix 1

Table of AICc values.

|  | Model* | number of<br>parameters | QAICc | Deviance | c-hat |
| --- | --- | --- | --- | --- | --- |
| Gulf of Lion | 1 | 9 | 983.89 | 126.80 | 1.39 |
|  | 2 | 3 | 1002.54 | 157.67 | 1.39 |
|  | 3 | 11 | 987.54 | 126.38 | 1.39 |
|  | 4 | 5 | 1006.58 | 157.67 | 1.39 |
| Riviera | 1 | 9 | 211.15 | 68.91 | 0.87 |
|  | 2 | 3 | 254.54 | 125.70 | 0.87 |
|  | 3 | 11 | 214.72 | 67.82 | 0.87 |
|  | 4 | 5 | 258.83 | 125.70 | 0.87 |
| Continental coast | 1 | 9 | 1130.85 | 129.72 | 1.60 |
|  | 2 | 3 | 1154.71 | 165.75 | 1.60 |
|  | 3 | 11 | 1132.26 | 127.03 | 1.60 |
|  | 4 | 5 | 1156.45 | 163.45 | 1.60 |

\* Models were built as follows:
Model 1 considers two age classes in survival and season-dependent recapture probabilities
Model 2 considers two age classes in survival and constant recapture probabilities
Model 3 considers two age classes in survival, heterogeneous and season-dependent
recapture probabilities
Model 4 considers two age classes in survival and heterogeneous recapture probabilities

